# Construction and Operation of an Affordable Laboratory Photobioreactor System for Simultaneous Cultivation of up to 12 Independent 1 L Cyanobacterial Cultures

**DOI:** 10.1101/153023

**Authors:** Ryan L. Clark, Laura L. McGinley, Daniel F. Quevedo, Thatcher W. Root, Brian F. Pfleger

## Introduction

A large field of research has developed for the study of cyanobacteria for photosynthetic chemical production [1]. Understanding the bioenergetics of growth and product secretion of cyanobacteria requires experiments in controlled environments with volumes large enough to allow sampling over time without removing most of the culture medium. The cost of commercial laboratory photobioreactors makes these experiments inaccessible to many researchers and limits experimental throughput. In this work, we designed and constructed a system of 12 independent, sterilizable cyanobacterial photobioreactors for cultures up to 1 L in volume for a cost less than a single commercial photobioreactor (Figure 1, Table 1). This system allows gas mixing to a desired CO_2_ concentration for transfer to the culture medium, discrete modulation of light intensity through addition/removal of fluorescent tubes, temperature control in the range of ambient temperature to 45°C, and simple sterile sampling for monitoring throughout long-term growth experiments. In the following sections, we will describe the assembly and operation of this photobioreactor system.

**Figure 1.**
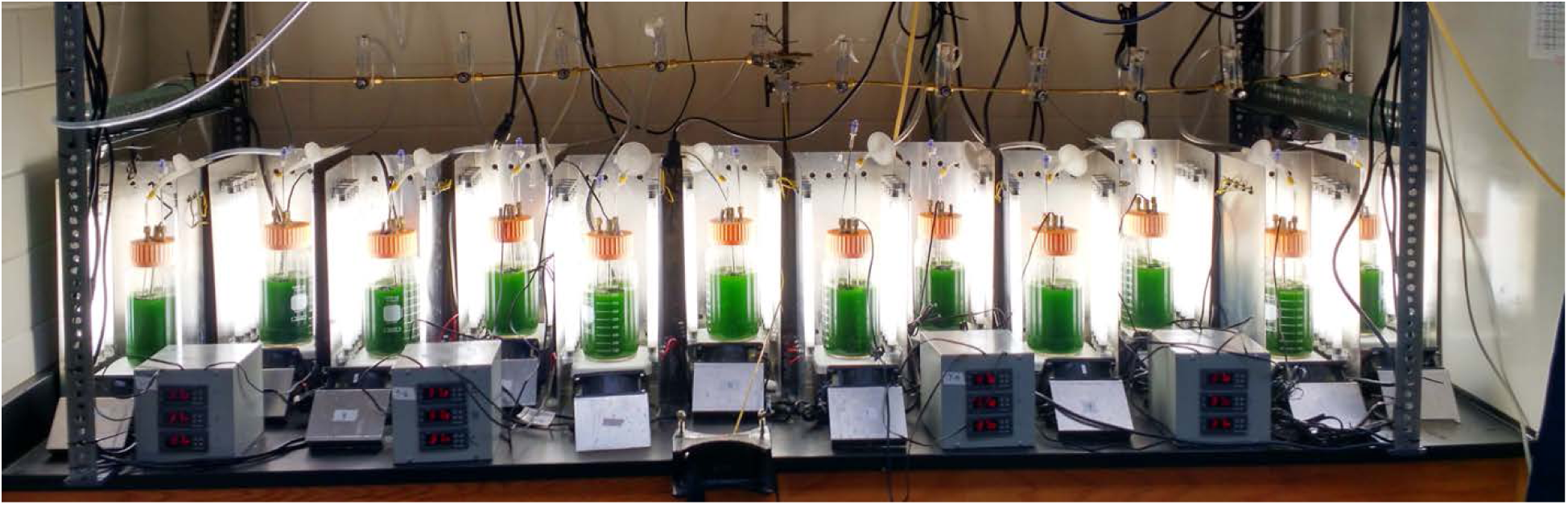
Photobioreactor System in Operation. This 12-reactor system was used to simultaneously grow cyanobacteria under varying conditions. Gas delivery could have the same composition for all 12 reactors or one composition for the first six (from the left) and a different composition for the second six. Temperature control systems were combined in sets of three, but temperature set points were independent for each reactor.

**Table 1.**
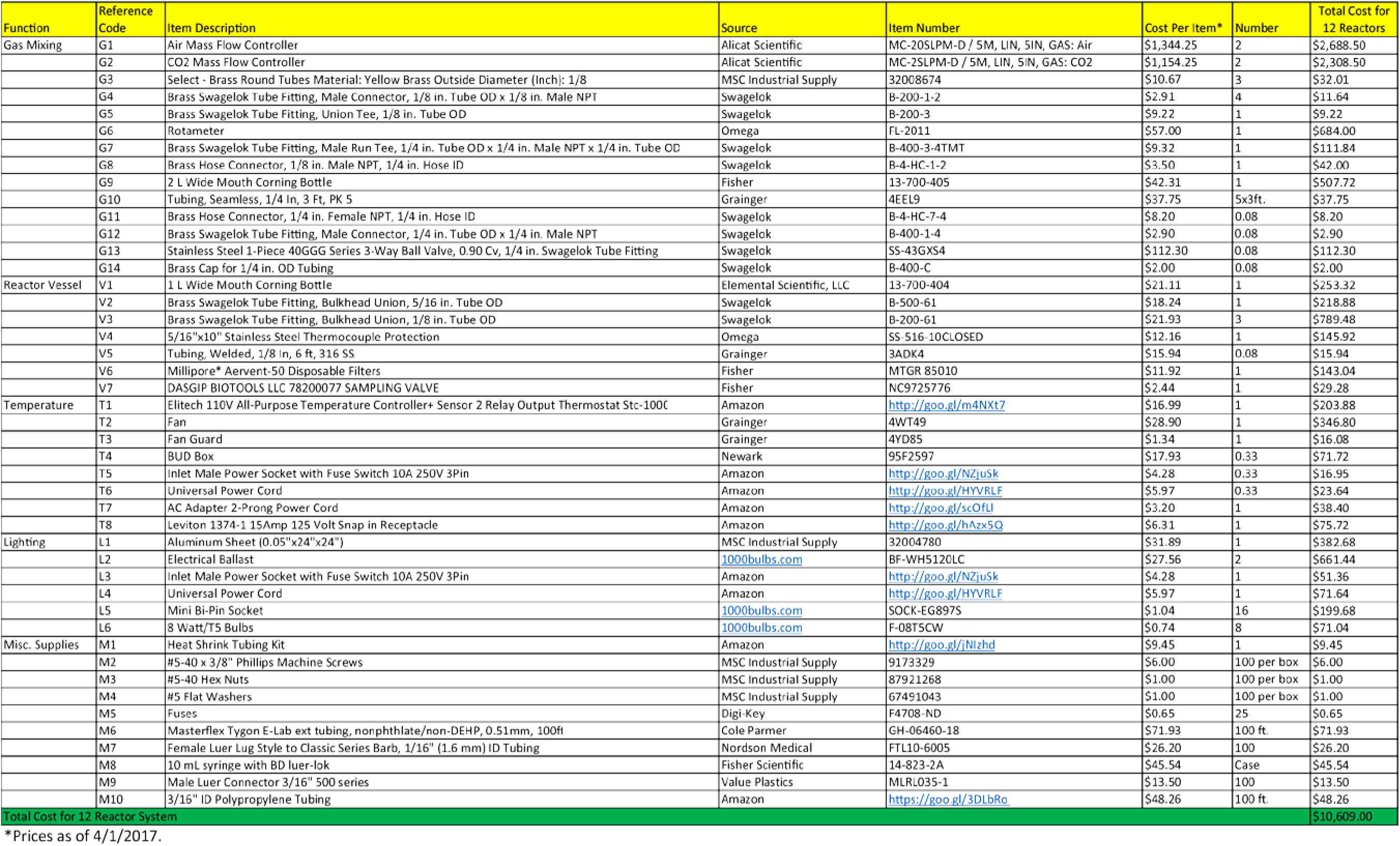
Parts List

## Photobioreactor System Assembly

### Reactor Vessel

1 L Wide Mouth Corning Bottles (V1) were used for the reactor vessel as they are a standardly available and cost-effective item that provided the desired working volume and substantial space for instrument installation in the cap. As shown in Figure 2, holes were drilled in the plastic cap of the vessel to allow installation of bulkhead fittings for (1) a sample port, (2) gas delivery, (3) off-gas exit, and (4) thermocouple protection.

**Figure 2.**
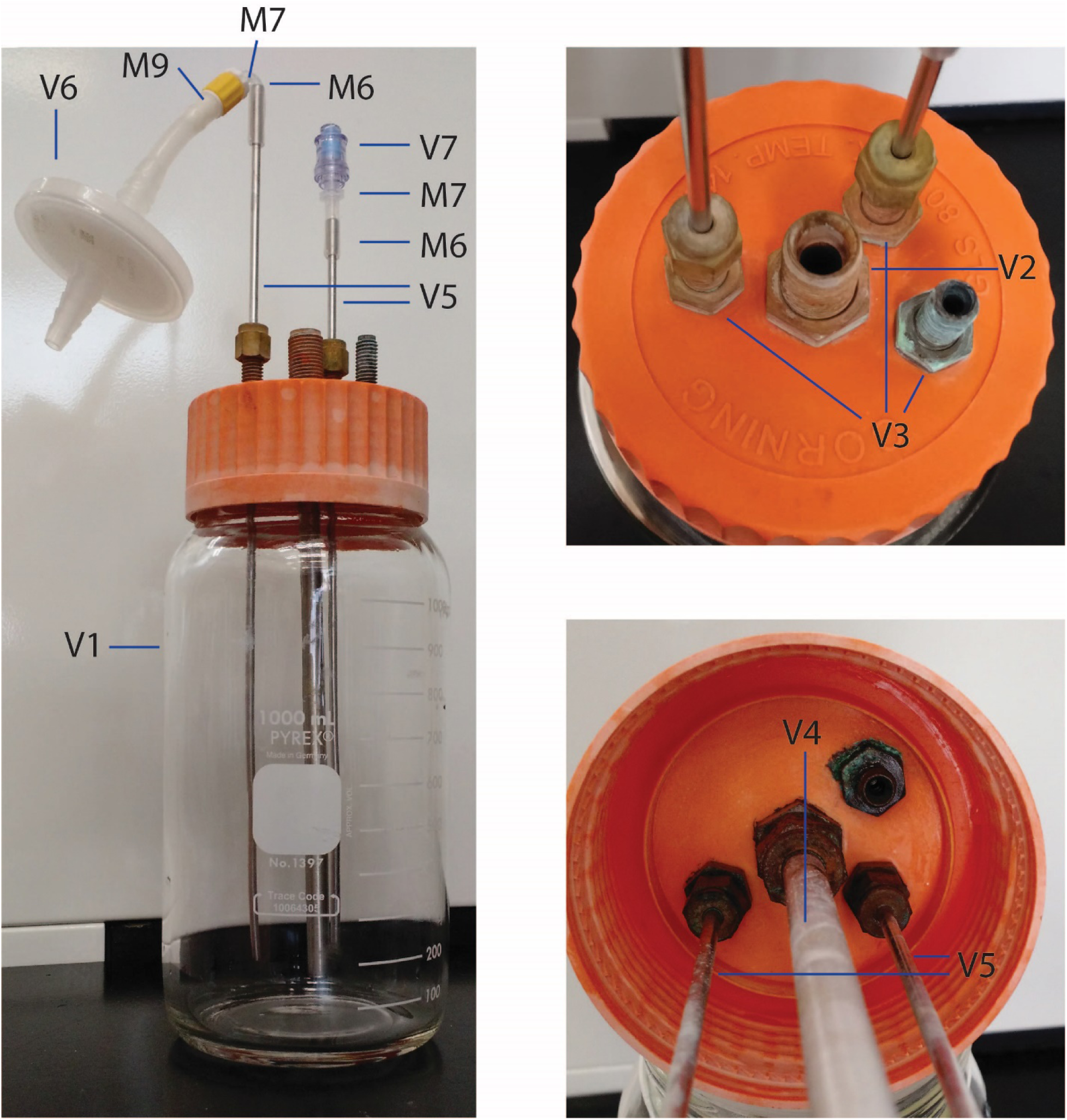
Reactor Vessel. (Left) Image of fully assembled reactor vessel. (Right Top) Top down view of reactor cap shows how instrumentation was installed on the top of the cap. (Right Bottom) Bottom up view of reactor cap shows how instrumentation was installed on the bottom of the cap. All labels reference the parts shown in Table 1.

The sample port used a Brass Swagelok Bulkhead Union for 1/8 inch Tube OD (V3). A short piece of 1/8 inch stainless steel tubing (V5) was attached to the Swagelok fitting of the bulkhead union (V3) on the outside of the cap which was connected to a short length of 0.51mm Masterflex Tygon E-Lab tubing (M6). To this tubing a female Luer lock to 1/16 inch barb connector (M7) was attached which was connected to a DASGIP Sampling Valve (V7). A 7.25 inch piece of 1/8 inch stainless steel tubing (V5) was attached to the Swagelok fitting of the bulkhead union (V3) on the inside of the cap. The resulting sample port allowed for online sampling of culture from near the bottom of the reactor using a Luer lock syringe.

The gas delivery port also used a Brass Swagelok Bulkhead Union for 1/8 inch Tube OD (V3). A short piece of 1/8 inch stainless steel tubing (V5) was attached to the Swagelok fitting of the bulhead union (V3) on the outside of the cap which was then connected to a short length of 0.51mm Masterflex Tygon E-Lab tubing (M6). To this tubing a female Luer lock to 1/16 inch barb connector (M7) was attached which was connected to a male Luer lock to 3/16 inch barb (M9). A short piece of 3/16 inch ID polypropylene tubing (M10) was attached to this 3/16 inch barb (M9) and subsequently to a Millipore Aervent-50 Disposable Filter (V6). This filter would eventually be attached to the gas delivery manifold discussed later. A 7.25 inch piece of 1/8 inch stainless steel tubing (V5) was attached to the Swagelok fitting of the bulkhead union (V3) on the inside of the cap. The resulting gas delivery port allowed delivery of sterile filtered gas to the bottom of the reactor vessel for CO_2_ delivery.

The off-gas exit port also used a Brass Swagelok Bulkhead Union for 1/8 inch Tube OD (V3). No experiments performed with this system have taken advantage of the off-gas exit port, but Swagelok fittings could be used to capture the off-gas for further analysis if desired.

The thermocouple protection port used a Brass Swagelok Bulkhead Union for 5/16 inch Tube OD (V2). A 5/16 inch stainless steel thermocouple protection piece (V4) was cut to a length of 7.25 inches and attached to the Swagelok fitting of the bulkhead union (V2) on the inside of the cap. The resulting thermocouple protection port allowed a thermocouple to sample the reactor temperature without contacting the culture medium.

The resulting reactor vessel was suitable for cultivation of photosynthetic microorganisms with culture volumes of 500-1000 mL in conjunction with the light delivery, gas delivery, and temperature control systems described hereafter.

### Gas Delivery System

An overview of the gas delivery system used to deliver a gas phase with a desired CO_2_ concentration is shown in Figure 3. Two Alitech Mass Flow Controllers (G1, G2) were used to mix ambient air and industrial CO_2_ to the desired partial pressure of CO_2_. The ambient air was provided by a house compressed air line fitted with a regulator to achieve the appropriate pressure. Industrial grade CO_2_ was provided by a gas cylinder fitted with a regulator to achieve the appropriate back pressure. All plumbing for the mixing system used brass tubing with 1/8 inch OD (G3) and brass Swagelok tube fittings. A piece of 0.51mm MasterFlex Tygon E-Lab tubing (M6) was attached to the end of the brass tubing extending from the point of mixing and attached to another piece of brass tubing entering the water bubbler to hydrate the gas phase before entering the reactor.

**Figure 3.**
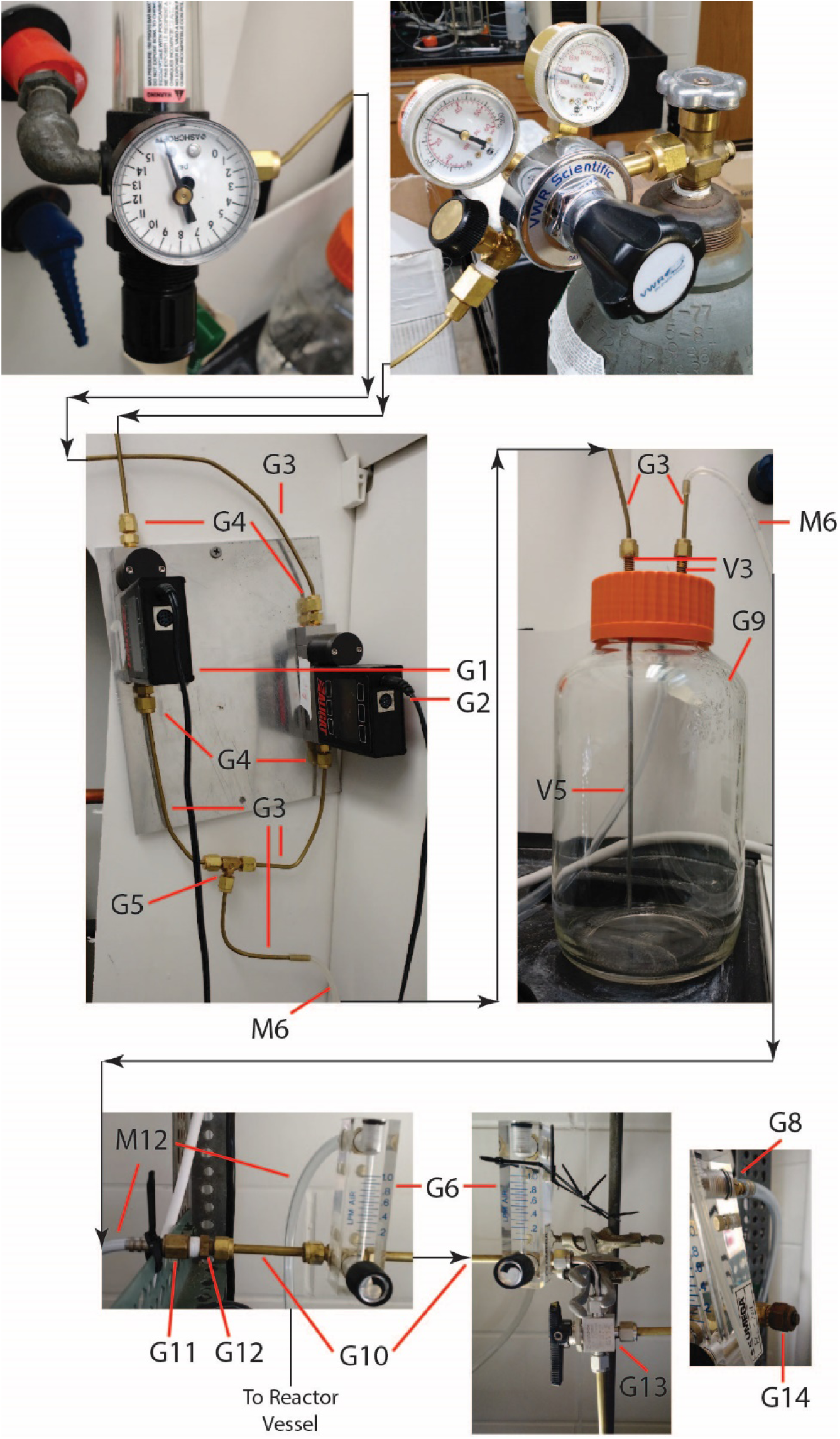
Gas Mixing System. (Top Left) Ambient air was fed from house air fitted with a regulator to achieve desired pressure. (Top Right) Industrial grade CO2 was fed from a gas cylinder fitted with a regulator to achieve desired pressure. (Middle Left) The flow rate of gases was controlled using mass flow controllers and mixed by combining the output flows. (Middle Right) The mixed gas stream was fed through a water bubbler to hydrate the air (shown without water). (Bottom Left) The hydrated gas phase was fed into a gas manifold for delivery to individual reactor vessels using rotameters to control gas flow rate. All labels reference the parts shown in Table 1.

The water bubbler was made using a 2 L Wide Mouth Corning Bottle (G9) with a cap machined to hold two Brass Swagelok Bulkhead Unions for 1/8 inch tube OD (V3), one for the gas inlet and one for gas outlet. A small piece of 1/8 inch OD stainless steel tubing (V5) was attached to the outside Swagelok fitting of each bulkhead union (V3) and a 7.25 inch piece of 1/8 inch OD stainless steel tubing (V5) was attached to the inside Swagelok fitting of the bulkhead union for gas inlet (V3) to submerge the inlet opening. This bubbler was filled approximately ¾ full with distilled water. A piece of 0.51mm MasterFlex Tygon E-Lab tubing (M6) was attached to the outlet. To this tubing a female Luer lock to 1/16 inch barb connector (M7) was attached which was connected to a male Luer lock to 3/16 inch barb (M9). A piece of 3/16 inch ID polypropylene tubing (M10) was attached to this 3/16 inch barb and subsequently to the gas delivery manifold. Two of these gas delivery systems were built to allow for simultaneous experimentation with different gas phases.

The gas delivery system consisted of 12 rotameters (G6) connected in parallel with ¼ inch OD brass tubing (G10) and brass Swagelok tube fittings. A three-way valve (G13) was placed after the sixth rotameter to allow two operating modes: one in which all 12 rotameters are fed from one gas mixing system and a second in which two different gas phases could feed the first six and the second six rotameters. In the first case, the 3/16 inch ID tubing (M10) from the first gas mixing system was attached to the manifold upstream of all 12 rotameters. In the second case, the 3/16 inch ID tubing (M10) from the second gas mixing system was attached to the manifold at the three-way valve (G13). Each rotameter had a brass hose connecter for 1/4 inch hose ID (G8) which was used to deliver gas to the inlet of each reactor. Flow rates for individual reactors were adjusted using the rotameter valve.

### Lighting System

The lighting system for the photobioreactors fulfilled two roles: providing light to the culture and providing heat to the system. The temperature control system described in the subsequent section provided cooling to offset the heat provided by the fluorescent tubes and maintain the desired temperature set point. An image of the lighting system is shown in Figure 4.

**Figure 4.**
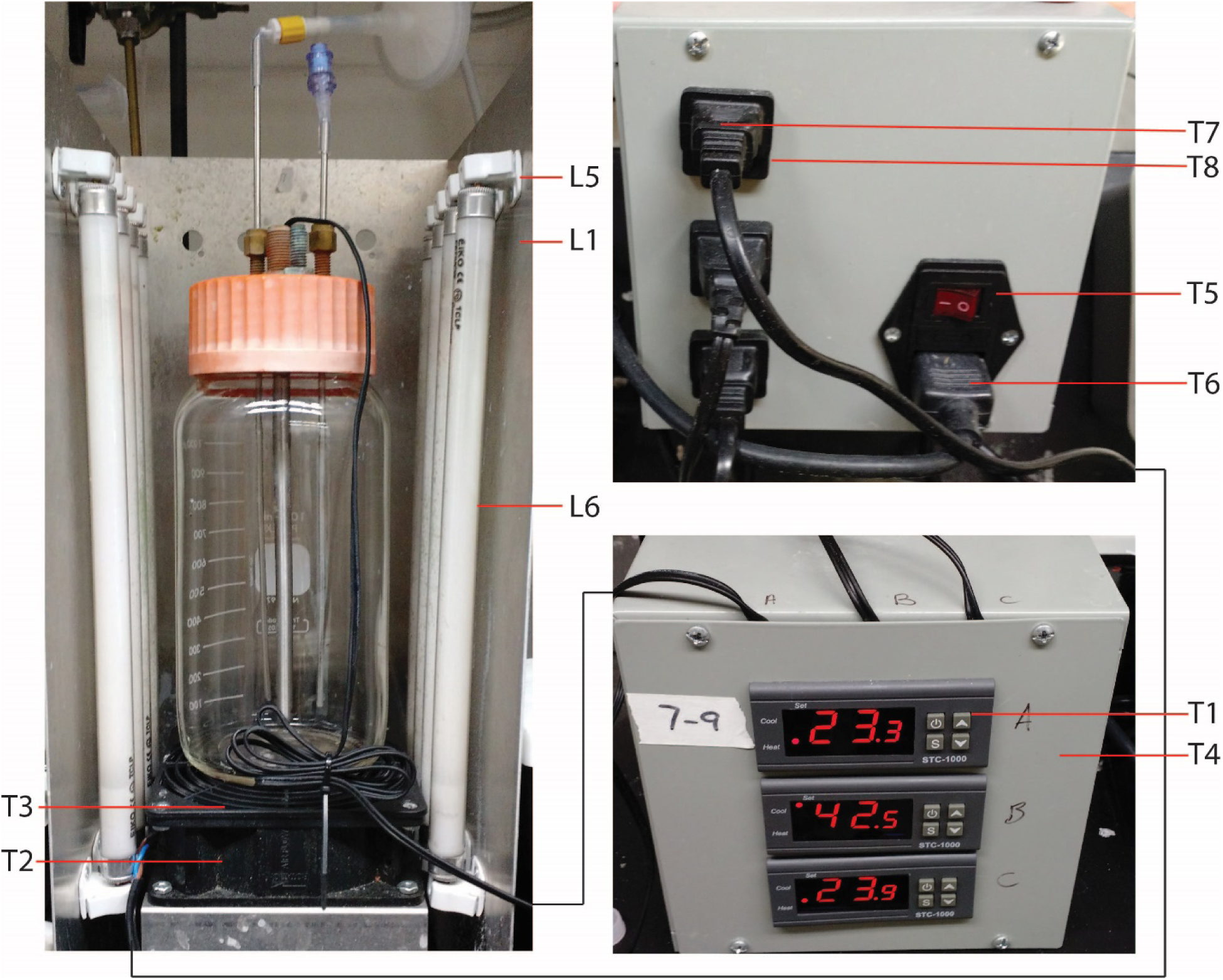
Lighting and Temperature Control System. (Left) The reactor vessel rested on top of the fan assembly (Figure 7) and light was provided from the sides of the reactor by the fluorescent light assembly (Figure 5). (Bottom Right) Front view of the temperature control assembly which used the temperature probe as an input. (Top Right) Back view of the temperature control assembly which turned the fan on or off per the control parameters of the temperature controller. The power input on the back of temperature control assembly provided power for both the controllers and the fans. All labels reference the parts shown in Table 1.

An overview of the assembly of the lighting system is shown in Figure 5. A piece of sheet metal (L1) was sheared to the appropriate size and holes were cut using a sheet metal punch. The sheet was then bent at a 90° angle using a bender brake. Each mini bi-pin socket (L5) had a hole drilled in its base using a drill press. This hole was used to affix the sockets to the inside of the sheet metal fixture using a machine screw (M2), washer (M4), and nut (M3). Wires from the ballasts (L2) were then inserted into the corresponding sockets to provide power to the fluorescent bulbs per the manufacturer’s instructions. Two ballasts were required for each lighting system, with each ballast powering 4 fluorescent tubes. Power was provided to the ballasts from a power socket with a fuse switch (L3) to allow the lights to be turned on and off. To simplify wiring, lighting systems were grouped into sets of three with one power socket with switch controlling all three lighting systems.

**Figure 5.**
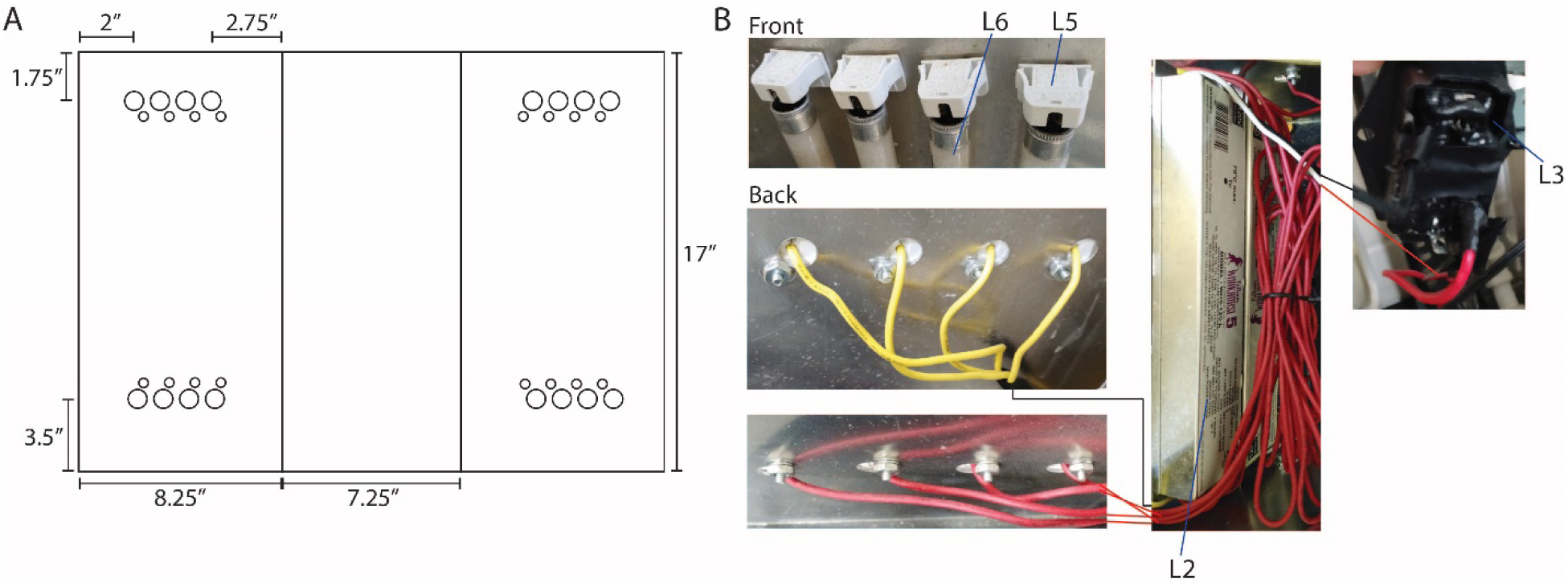
Lighting System Assembly. (A) The sheet metal (L1) was cut to the size shown here and holes were punched for fastening and wire access to the mini bi-pin sockets for the fluorescent bulbs (L5). The sheet metal was folded at a ninety-degree angle on the two indicated lines to form the three-sided shape allowing the assembly to stand up. (B) (Top Left) L5 were installed on the inside of the sheet metal casing at all the locations indicated in (A) for a total of 16. These held the fluorescent tubes in place and provided power from the ballasts. (Middle Left) the sockets on the top of each side were wired to the yellow leads from the ballast (4 bulbs per ballast). (Middle Bottom) the sockets on the bottom of each side were wire to the red leads from the ballast (4 bulbs per ballast). (Center) The ballasts were fixed to the back of the sheet metal assembly. (Right) The black and white power supply leads from the ballasts were wired to a power socket for control of power supply. All labels reference the parts shown in Table 1.

Experiments in this work were performed using 4100K cool white fluorescent tubes (L6). This lighting system will work with any F8T5/CW 8-watt fluorescent tubes if a different light quality is desired. Light intensity was modulated in a discrete manner by changing the number of fluorescent bulbs installed in the system as shown in Figure 6. Luminous flux was determined with a Traceable Dual-Range Light Meter (Fisher Scientific).

**Figure 6.**
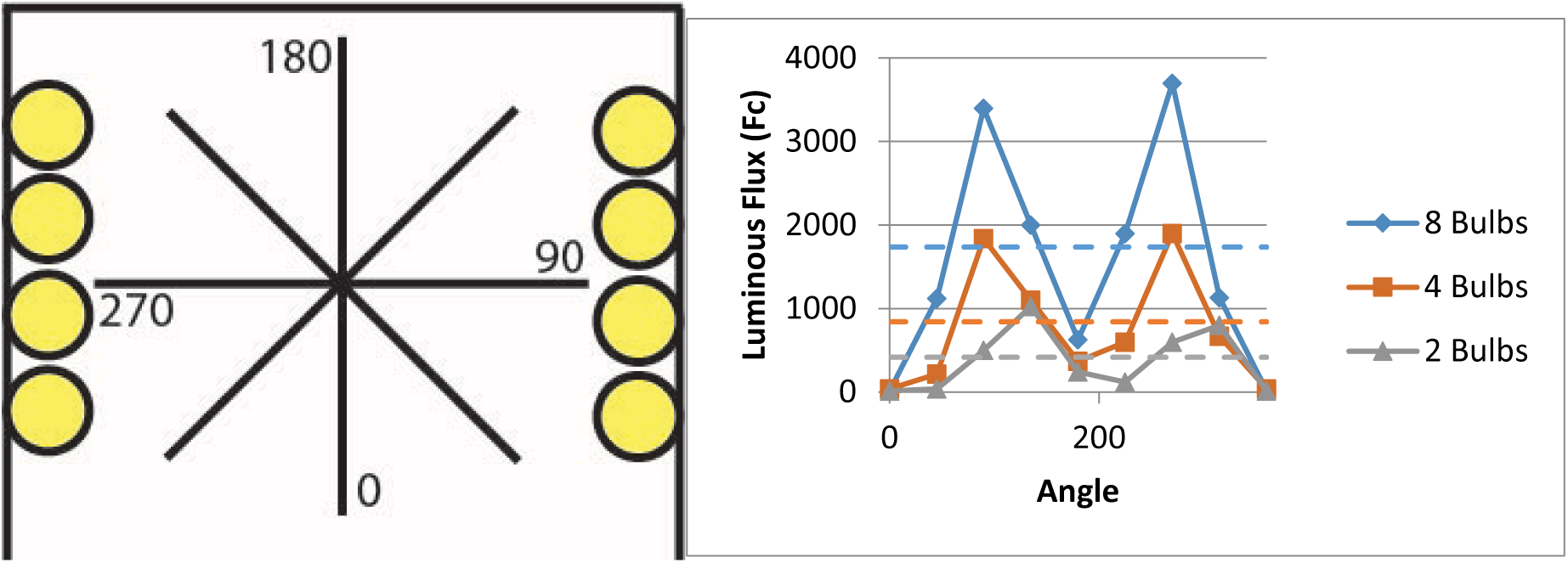
(Left) Top down schematic of lighting system. 4 bulb set up had second and fourth bulb remove on the left and first and third bulb remove on the right. 2 bulb set up had only top bulb on the right and only the bottom bulb on the left. Numbers represent the angles at which luminous flux was measured. (Right) Luminous flux as a function of angle for each lighting setup. The dashed line represents the average luminous flux for a given number of bulbs.

### Temperature Control

As mentioned above, heat was provided to the system by the fluorescent tubes. Without cooling and with all 8 bulbs installed in a lighting system, the temperature of liquid in the reactor vessel rose to over 40°C. To maintain cultures at a constant temperature, a temperature control system provided cooling based on temperature measurements as shown in Figure 4. The thermocouple inserted into the stainless-steel thermocouple protector on the reactor vessel detected the temperature and the temperature controller output power to the corresponding fan to cool the reactor to the desired set point. The temperature set point could be achieved for any temperature between ambient room temperature and 40°C. The temperature controller and fan were wired per the manufacturer’s instructions as shown in Figure 7.

**Figure 7.**
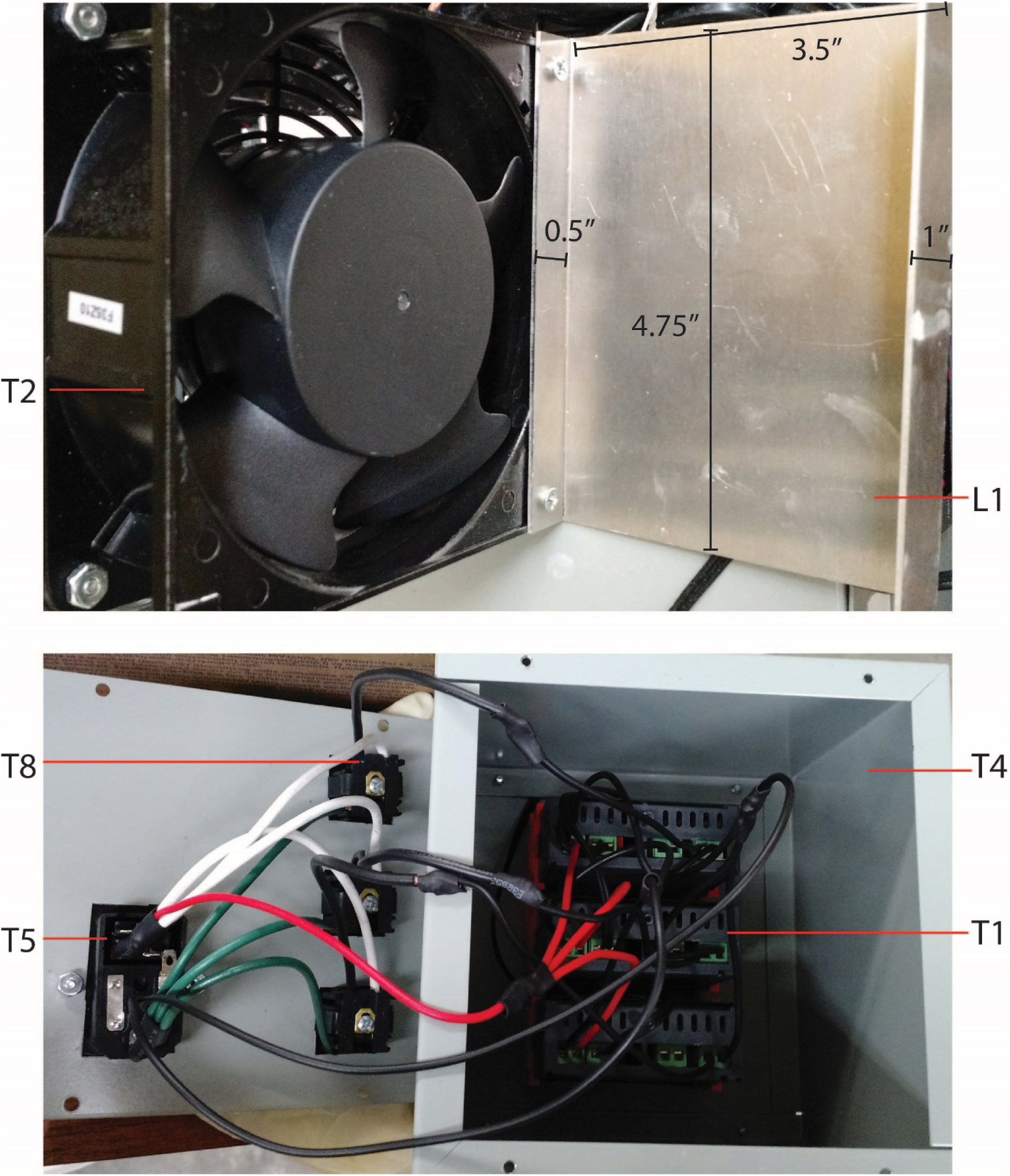
Temperature Control System Assembly. (Top) Sheet metal supports for the fan were machined and fixed to the fan as shown allowing the fan to sit upright and act as a stand for the reactor vessel (one support removed for image). (Bottom) Temperature controllers were wired per the manufacturer’s instructions with the power supply coming from the power socket (T5) and the output for cooling leading to the snap in receptacles (T8). All labels reference the parts shown in Table 1.

## Example Operating Procedure

### Media Preparation

Media A was used in the photobioreactors for cultivation of *Synechococcus* sp. PCC7002 (components shown in Table 2, adapted from Stevens, et al.[2]). All components were added in the order listed to a 50 L carboy, and the solution was mixed between additions by shaking. Media was made in one large batch to be distributed to the photobioreactors for autoclaving. Sodium chloride and magnesium sulfate were added as solids and each dissolved completely before the addition of the next component. All other components were added from 100x stock solutions, with the trace elements combined into a single 100x stock. To prepare this stock, ferric ammonium citrate, boric acid, and manganese chloride were added as solids, and the remaining components were added using stocks of the concentrations shown in Table 3. The trace elements stock was filter sterilized immediately after preparation.

**Table 2.** Target concentrations of all media components.

| Component | Concentration<br>(mM) |
| --- | --- |
| **NaCl** | 308 |
| **MgSO<sub>4</sub>·7H<sub>2</sub>O** | 20 |
| **Na<sub>2</sub>EDTA·2H<sub>2</sub>O** | 0.08 |
| **KCl** | 8.1 |
| **CaCl<sub>2</sub>·2H<sub>2</sub>O** | 2.5 |
| **NaNO<sub>3</sub>** | 12 |
| **KH<sub>2</sub>PO<sub>4</sub>** | 0.37 |
| **Tris** | 8.3 |
| **Trace elements** | (uM) |
| Ferric Ammonium Citrate | 30 |
| H<sub>3</sub>BO<sub>3</sub> | 554 |
| MnCl<sub>2</sub> 4H<sub>2</sub>O | 22 |
|  | (nM) |
| ZnCl<sub>2</sub> | 2310 |
| MoO<sub>3</sub> | 208 |
| CuSO<sub>4</sub>·5H<sub>2</sub>O | 12 |
| CoCl<sub>2</sub>·6H<sub>2</sub>O | 51 |

**Table 3.**
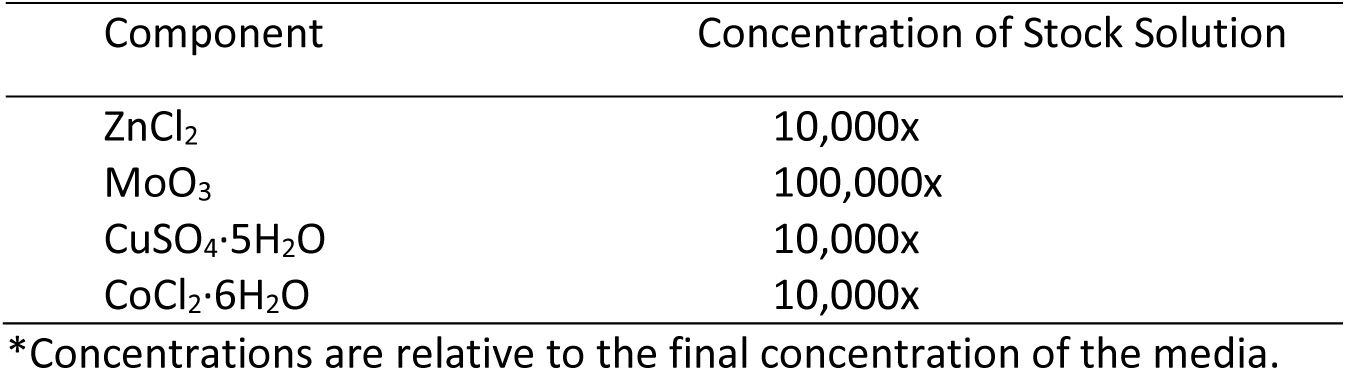
Concentrations of stock solutions of trace element stock ingredients

| Component | Concentration of Stock Solution |
| --- | --- |
| ZnCl<sub>2</sub> | 10,000x |
| MoO<sub>3</sub> | 100,000x |
| CuSO<sub>4</sub>·5H<sub>2</sub>O | 10,000x |
| CoCl<sub>2</sub>·6H<sub>2</sub>O | 10,000x |

\*Concentrations are relative to the final concentration of the media.

### Reactor Assembly and Sterilization

900 ml of media was used in each reactor. Immediately before media was measured and added to the reactors, it was mixed by shaking to suspend any small amount of precipitate that may have settled out. After the reactors were filled, the caps were loosely attached and wrapped in foil. They were then autoclaved for 40 minutes to sterilize the media and internal components. Foil remained over the gas outlet until the reactors were attached to the air delivery system to block any convection of nonsterile air into the reactor. B12 (typically via a 1000x stock solution) was added to achieve a concentration of 3 nM, and any necessary antibiotics and inducers were also added. The reactors were then inoculated to a target OD730 of 0.05 from a pre-culture of the organism to be studied.

### Sampling Protocol

Prior to sampling, bioreactor volumes were adjusted to account for evaporation losses by adding sterile water through the DASGIP sampling valve. Evaporation losses were typically about 10-20 ml per day and slightly variable among reactors. Sterile water was stored in a bottle fitted with a DASGIP sampling valve of the same type used on the bioreactor assembly. Both valves were sterilized with a 70% aqueous ethanol solution immediately before a syringe was used to transfer the water.

The syringe was then used to sample the reactor. During sampling, the syringe was filled and emptied several times while attached to the reactor before collecting the final sample to flush out the sample port. Sample volumes were typically 1 mL or less to measure cell density and pigmentation, although larger volumes were taken for more complex sample analysis. Samples of 1 ml or less were regarded as negligible to the overall reactor volume, but larger samples were accounted for when adjusting for evaporation losses the following day. The OD730 was measured to observe cell density over time. Figure 8 shows a representative growth curve for Synechococcus sp. PCC7002 grown in Media A in the photobioreactors.

**Figure 8.**
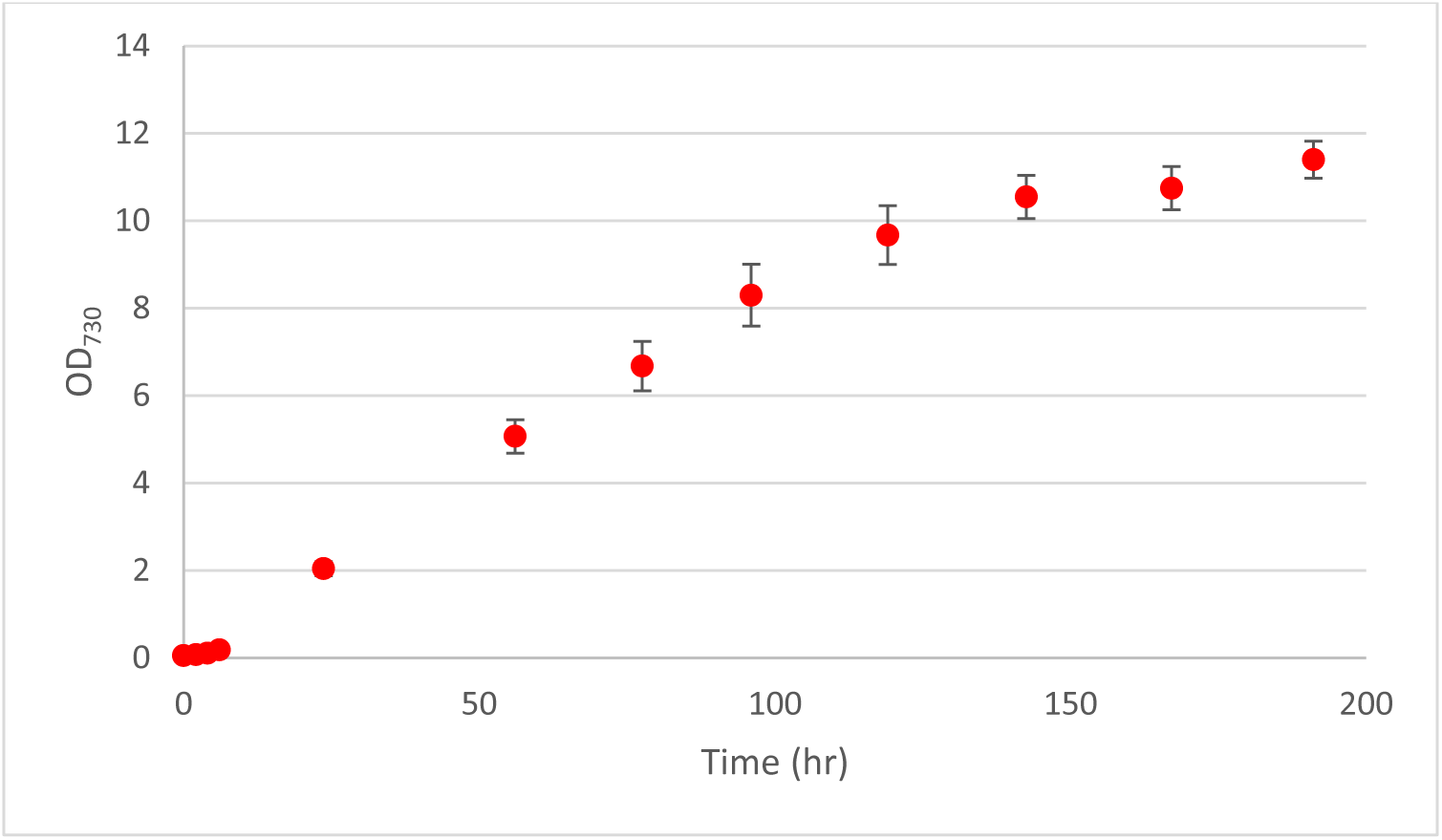
Representative growth curve for *Synechococcus* sp. PCC7002 in photobioreactors. 900 mL cultures of *Syenchococcus* sp. PCC7002 were grown in the photobioreactors bubbled with 10% CO_2_ and maintained at a temperature of 37°C by the temperature control system. Error bars are standard deviation of two biological replicates.

